# Tool use fine-tunes arm and tool maps

**DOI:** 10.1101/2024.06.06.597694

**Authors:** V.C. Peviani, L.N. Pfeifer, G. Risso, M. Bassolino, L.E. Miller

## Abstract

There is evidence that the sensorimotor system builds fine-grained spatial maps of the limbs based on somatosensory signals. Can a hand-held tool be mapped in space with a comparable spatial resolution? Do spatial maps change following tool use? In order to address these questions, we used a spatial mapping task on healthy participants to measure the accuracy and precision of spatial estimates pertaining to several locations on their arm and on a hand-held tool. To study spatial accuracy, we first fitted linear regressions with real location as predictor and estimated location as dependent variables. Intercepts and slopes, representing constant offset and estimation error, were compared between arm and tool, and before to after tool use. We further investigated changes induced by tool use in terms of variable error associated with spatial estimates, representing their precision. We found that the spatial maps for the arm and tool were comparably accurate, suggesting that holding the tool provides enough information to the sensorimotor system to map it in space. Further, using the tool fine-tuned the user’s spatial maps, increasing the precision of the tool map to a greater extent than their arm map. Furthermore, this increased precision is focal to specific tool locations, i.e., the tool tip, which may work as a spatial anchor following tool use. Our results demonstrate that tool users possess dynamic maps of tool space that are comparable to body space.

## Introduction

For perception and motor control, the brain relies on internal representations of the body that specify the configuration of the limbs in space—essential for interaction with physical objects in the environment (Medina & Coslett, 2010; Tamè et al., 2019). Previous research measuring spatial maps of limbs has shown that healthy participants can successfully map the spatial layout of their body parts, such as the hands (e.g., Longo & Haggard, 2010), legs (Stone et al., 2021), and arms (e.g., Bassolino et al., 2015; Galigani et al., 2020). These spatial estimates however are not free from biases (e.g., fingers are perceived as shorter: Longo & Haggard, 2010; Peviani & Bottini, 2020), which may partially arise from the probabilistic computations involving spatial priors (Peviani et al., 2024). Interestingly, some studies reported changes in the pattern of biases based on motor behavior (e.g., Canzoneri et al., 2013) and expertise (e.g., Coelho et al., 2019). In all, these data suggest that body maps are spatially defined and support sensorimotor behavior (Peviani et al., 2020; Peviani & Bottini, 2018).

The evolution of tools has further heightened human capabilities, acting as physical and functional extensions of our limbs (Cardinali et al., 2009; Maravita & Iriki, 2004; Martel et al., 2016; Miller et al., 2014), and ultimately extending somatosensory processing beyond the body (Giudice et al., 2013; Miller et al., 2018; Yamamoto & Kitazawa, 2001). The sensorimotor system quickly adapts to a hand-held tool, changing motor control policies (Itaguchi & Fukuzawa, 2014; Martel et al., 2016) and altering the user’s action-oriented body representations (Cardinali et al., 2009, 2011; Miller et al., 2017). Tools can also function as *sensory* extensions of the body (Giudice et al., 2013). Humans can accurately localize touch when applied on the surface of an unseen hand-held tool (Miller et al., 2018), including rods that are six-meters in length (Miller et al., 2023).The brain may repurpose computations to control and perceive a limb to control and perceive a tool (Head & Holmes, 1911; Martel et al., 2016; Miller, Fabio, et al., 2023).

The above findings suggest that tool users can infer the spatial properties of a rod when striking an object. This is likely because they have internal models of the rod’s dynamics (e.g., torque, muscle stretch, vibrations) that allow for accurate control and sensing (Imamizu et al., 2000, 2003; Kilteni & Ehrsson, 2017; Miller et al., 2018). Humans can also sense the rudimentary geometry of a rod (e.g., length) just by holding it, presumably by tuning into how it alters proprioceptive feedback of the limb (Debats, Van De Langenberg, et al., 2010; Solomon & Burton, 1989; Burton & Turvey, 1990). However, it is unknown if the somatosensory system also builds up a fine-grained spatial map of a tool as it does for the body, and to what extent the two are comparable. It is further unknown whether using a tool alters the spatial map of the tool as it does for maps of the body (e.g., Canzoneri et al., 2013; Cardinali et al., 2009, 2011; Galigani et al., 2020; Martel et al., 2016; Miller et al., 2014, 2017). In the present study, we aimed to fill these gaps.

To test these ideas, we adapted a spatial localization task designed by Longo & Haggard (2010) to measure fine-grained spatial perception of an unseen tool and of the arm holding it. Our novel design allowed us to map perceptual estimates of several locations on the forearm and hand-held tool *together*, and investigate if and how they change after tool use (10 minutes of visuo-motor interaction with the tool during an object-manipulation task). We first quantified the accuracy of the spatial tool and arm maps and how they were altered by tool exposure, by analyzing patterns of systematic errors in perceptual estimates of locations. We then quantified the spatial precision of arm and tool maps by analyzing the variable errors and their changes after tool use. This rather unexplored variable represents the spatial tuning of the maps, which may be decreased at anchor spatial locations such as limb joints and endpoints (Miller et al., 2022). To anticipate the results, we found that the spatial maps of the user’s arm and tool were comparably fine-grained. Further, using the tool fine-tuned the user’s spatial maps, altering the precision of the tool map to a greater extent than their arm map. In total, our results demonstrate that tool users possess dynamic maps of tool space that are comparable to body space.

## Methods

### Participants

Twenty participants (13 females) took part in the experiment. All participants were right-handed, had normal or corrected-to-normal vision, were 18 years of age or older (age range: 19-27, mean±std 22.69 ± 2.39 years), were English speaking, and had no motor impairments. The experiment was approved by the Ethics Committee of the Faculty of Social Sciences, Radboud University, Nijmegen (NL). All participants gave written consent before their participation in the experiment. Participants were recruited through the SONA system of Radboud University, and received as compensation course credits or a 10 Euro voucher.

### Task and procedure

Participants performed a spatial mapping task (e.g., Longo & Haggard, 2010; Peviani & Bottini, 2018) that simultaneously measured spatial maps of the forearm and hand-held tool. The experiment followed a pre-post design, in which measurements of the spatial map of the forearm and tool were carried out before and after a tool use task that required sorting and manipulating objects.

#### Spatial mapping task

Participants were seated in front of a screen which was placed horizontally on a table right above their unseen right arm and hand-held tool (**Figure 1A**), a 40-cm mechanical grabber (**Figure 1B**). When held, the tool had a functional length of about 34 cm. The elbow of each participant was fixed at the same pre-defined location on the table and relative to the screen. After the participant firmly grasped the tool below the screen, the tool was fixed and aligned with the forearm. The forearm and tool were both aligned with the screen’s longitudinal axis. A second screen was placed vertically in front of the participant to display a schematic drawing of an upper limb holding a tool, which was used to communicate the localization instructions for each trial.

**Figure 1.**
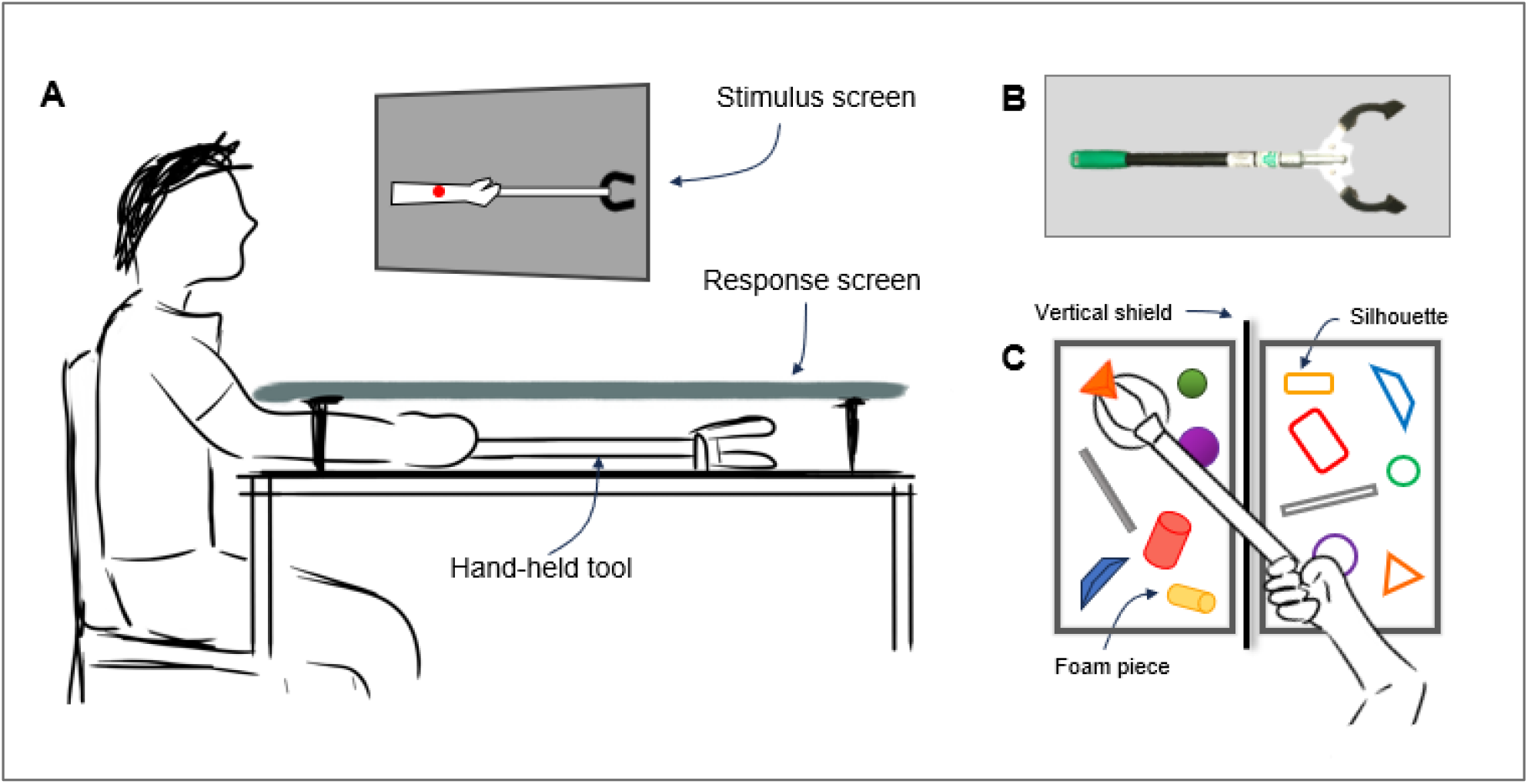
Experimental set-up and task. **A**. Arm and tool mapping task: Participants firmly held the tool with their right hand, under the response screen. A stimulus screen placed in front of the participant displayed one out of twelve target locations (six on the forearm, six on the tool) as a red dot. Participants were asked to estimate the position of this location in space by moving a cursor displayed on the response screen. The task was repeated after the tool use task. **B**. Forty-cm mechanical grabber used in the experiment. **C**. Tool use task: participants were instructed to grab different foam pieces from a box and move them over a shield to another box, by matching each foam piece with the right silhouette. Once all pieces were transferred in the correct positions, participants were instructed to move them back to the original box. This was repeated until 10 minutes elapsed.

The task for the participants was to localize a landmark on either their forearm or the hand-held tool. There were six equally-spaced landmarks that divided up the space (0–100%, by 20%) of the forearm (elbow-to-wrist) and rod (base-to-tip). Note that the location of arm landmarks, computed relative to the elbow, differed across participants as a function of the physical length of their forearm (mean±sem: 25.02 ± 0.41 cm). The location of the tool landmarks was calculated relative to the tool base, whose exact distance from the elbow was measured using measuring tape once the tool was fixed in position, before the mapping task. At the start of each trial, the landmark to be located was indicated on the arm+tool drawing with a red dot within the arm+tool space. Participants then moved a red cursor (controlled with a trackball by the left hand) above their physical (right) arm to the corresponding location in physical space and indicated with a mouse-click when they had reached the location of the landmark. The next trial then started. No feedback on the accuracy of the judgement was given to the participants. The cursor’s movement was constrained to a single degree of freedom, along a proximo-distal line aligned with the forearm.

Before starting the mapping task, participants could briefly see the tool while they were given instructions: no manipulation was allowed–besides holding the tool below the screen–before the tool use task. The mapping task was repeated once before and once after the object manipulation via tool use. Participants performed ten judgements per landmark, for a total of 120 judgements (randomized order) before and 120 after tool use. The task lasted on average 18.9 minutes (sd: 6 minutes) in the pre tool use phase, and 15.5 minutes (sd: 3.8 minutes) in the post tool use phase. Task control was implemented in MATLAB (R2022b).

#### Tool use task

Participants used the 40-cm mechanical grabber to grasp and move foam pieces from one box over one shield to another box, which was placed directly in front of participants. Each foam piece had a different shape, and participants were instructed to move and place each piece onto the corresponding silhouette drawn in the box **(Figure 1C)**. Participants had full visual feedback of their arm and tool. The tool use paradigm lasted 10 minutes. On average, participants grabbed, placed, and aligned 74 pieces (sd: 23).

#### Analysis

For each trial, actual locations and responses in pixel coordinates were extracted and transformed into centimeters (cm). First, we aimed to study accuracy of arm and tool maps. To do so, we fit four linear regression models to participant’s mean judgements in cm for arm (pre and post tool use) and tool (pre and post tool use), with actual locations as predictors. This allowed us to extract intercepts and slopes that serve as proxy of the arm and tool spatial maps. Intercept values represent general spatial biases in localization responses, with intercepts equaling zero representing no bias. In our set-up, intercepts less than zero indicate a proximal bias (i.e., spatial locations are perceived more towards the torso) whereas intercepts greater than zero a distal bias, which would manifest consistently across all spatial locations. Slope values indicate how spatial perception changed across consecutive spatial locations: when slopes equal one, estimated arm or tool length is accurate, whereas slopes less or greater than one indicate length underestimation or overestimation, respectively.

We then investigated the accuracy of arm and tool spatial maps *before* tool use by running one-sample tests on the pre tool use intercepts (against zero, i.e., no constant bias), and slopes (against one, i.e., no change in bias across locations). After, we evaluated differences in the accuracy of arm and tool maps, as well as changes from before to after tool use by inputting intercepts and slopes into two 2×2 repeated measures ANOVAs with two-level factors: surface (arm, tool) and time (pre, post). Second, we aimed to study the spatial precision of arm and tool maps; to this aim we calculated the variable error (i.e., standard deviation of the responses) for each of the twelve target locations, pre and post tool use. We used a 6×2 repeated measured ANOVA separately for arm and tool to investigate the change in variable error across the six landmarks (six-level factor: landmark position) from before to after tool use (two-level factor: pre, post).

ANOVAs were performed separately for each surface because there is no direct correspondence between the six landmarks on the arm and the tool. Therefore, we took another approach to compare how much tool use changed the variable errors for the tool and arm. To do so, we computed the Euclidean distance between the pre and post tool use variable error for each participant and surface (arm, tool). This quantity reflects how much tool use changed the spatial precision of each representation. Then, we compared the pattern of change between arm and tool using a paired t-test.

## Results

We first characterized the baseline accuracy of the arm and tool maps. The linear models fitted on mean perceived locations yielded very good fits (**Figure 2A**), with all R^2^ > 0.90 (arm, pre: R^2^ = 0.96; arm, post: R^2^ = 0.99; tool, pre: R^2^ = 0.96; tool, post: R^2^ = 0.95). We detected a significant general bias for arm and tool data *before* tool use (**Figure 2B**, top), with intercepts significantly lower than zero for the arm (mean±sem: -2.28 ± 0.57, t(19) = -3.86, p = .001) and tool maps (mean±sem - 4.41 ± 1.36, t(19) = -3.16, p = .005). Slopes were significantly lower than one for the arm (mean±sem: 0.87 ± 0.02, t(19) = -5.55, p <.001), while the difference was marginal for the tool (mean±sem: 0.91 ± 0.04, t(19) = -2.08, p = .051), **Figure 2B**, bottom.

**Figure 2.**
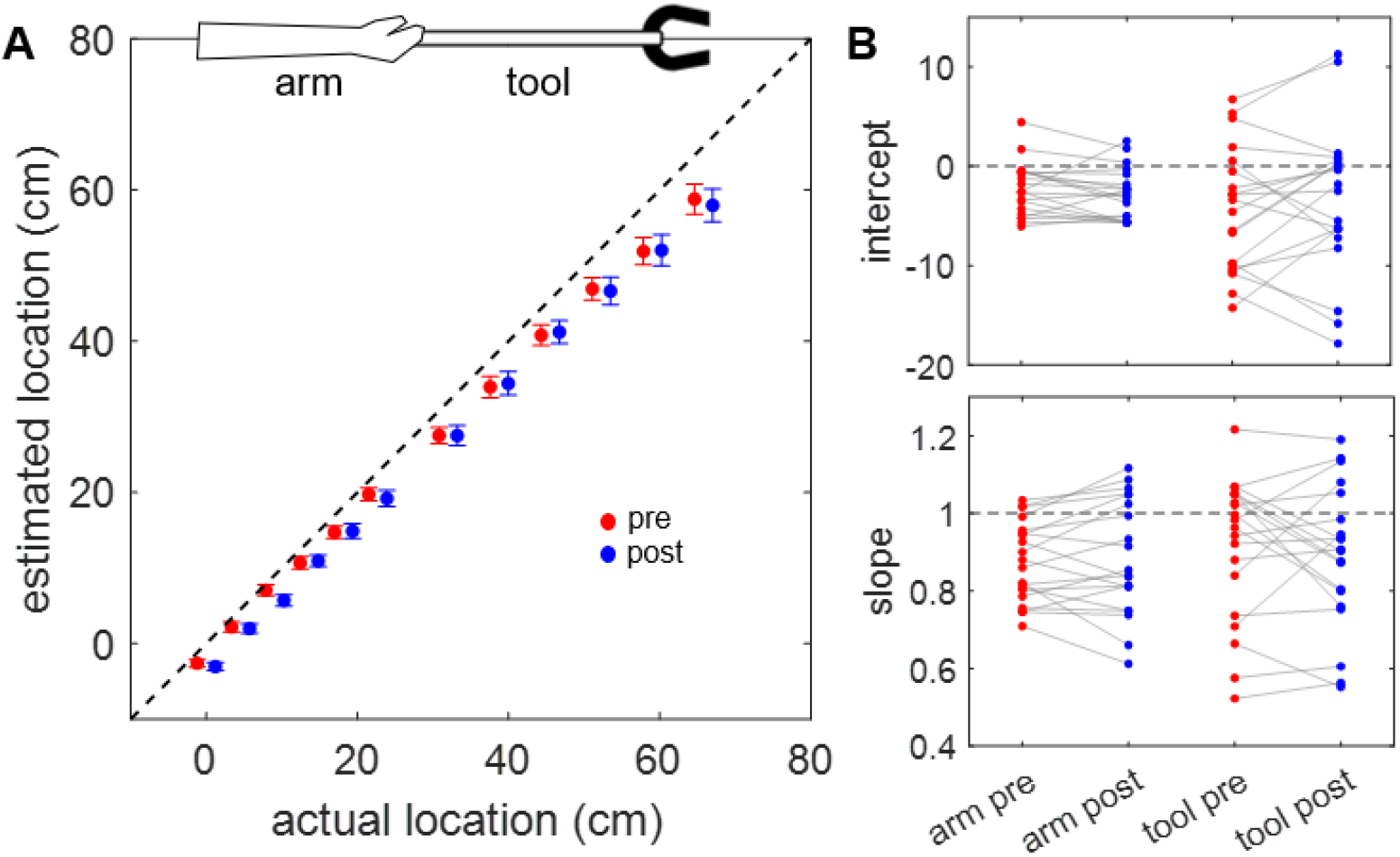
Accuracy of arm and tool mapping. **A**. The mean estimated locations of the twelve target positions (six for the arm, and six for the tool) are plotted against their actual locations in space. Error bars represent standard error of the mean. Red dots represent estimates before tool use, blue dots represent estimates after tool use. The dashed line is the identity line (i.e. slope equal to 1). Note that the pre and post dots have been artificially offset on the x-axis for ease of viewing. **B**. Intercepts and slopes estimated via linear regression are plotted for the arm and tool for each participant. The dashed lines represent no bias.

We next looked at the effect of tool use on the arm and tool maps. The 2×2 ANOVA for the intercept (**Figure 2B**, top) revealed no significant main effect of surface (F(1,19) = 0.98, p = .335, 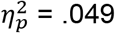), main effect of time (F(1,19) = 0.19, p = .667, 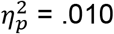), or an interaction (F(1,19) = 1.55, p = .230, 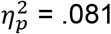). Indeed, the use-driven changes in the intercept were minimal for the arm (Δ(post-pre): - 0.51 ± 0.45) and tool (Δ(post-pre): 1.04 ± 1.14). The 2×2 ANOVA for the slope (**Figure 2B**, bottom) also revealed no significant main effect of surface (F(1,19) = 0.24, p = .635, 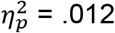), main effect of time (F(1,19) = 0.10, p = .749, 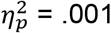), or an interaction (F(1,19) = 1.08, p = .310, 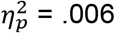). Indeed the use-driven changes in the slope were minimal for the arm (Δ(post-pre): 0.01 ± 0.08) and tool (Δ(post-pre): -0.02 ± 0.14. In general, these findings demonstrate (1) that the accuracy of the arm and tool maps were of a similar level of accuracy, and (2) that tool use did not alter the accuracy of either the arm or tool map.

We next investigated whether the precision of the spatial maps changed after tool use. To do so, we ran repeated-measures ANOVAs (2x6, with time and locations as factors) on the variable errors for the arm and tool separately. For both the arm and tool, we found a significant time-by-location interaction, demonstrating that using a tool differentially affected precision across the arm/tool surface (**Figure 3A**). In more detail, the interaction was significant for the arm (F(5,95) = 2.90, p = .018, 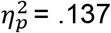) as well as for the tool (F(5,95) = 3.52, p = .006, 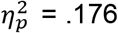). Each surface deviated in how tool use changed the pattern of variable errors. Post-hoc tests comparing variable errors measured before and after tool use for each location revealed for the arm lower variable errors close to the forearm midpoint (location 3 out of 12 with t(19) = 3.34, p = .018, bonferroni corrected), whereas for the tool variable errors were lower at the tool tip (location 12 out of 12 with t(19) = 3.89, p = .006, bonferroni corrected). Furthermore, for the arm main effects were also significant: main effect time, (F(1,19) = 9.45, p = .009, 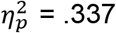); main effect location, F(5,95) = 19.67, p < .001, 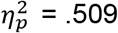. For the tool, we found a significant main effect of time (F(1,19) = 21.84, p = .006, 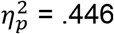), while the main effect of location was not significant (F(5,95) = 1.16, p = .331, 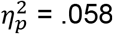).

**Figure 3.**
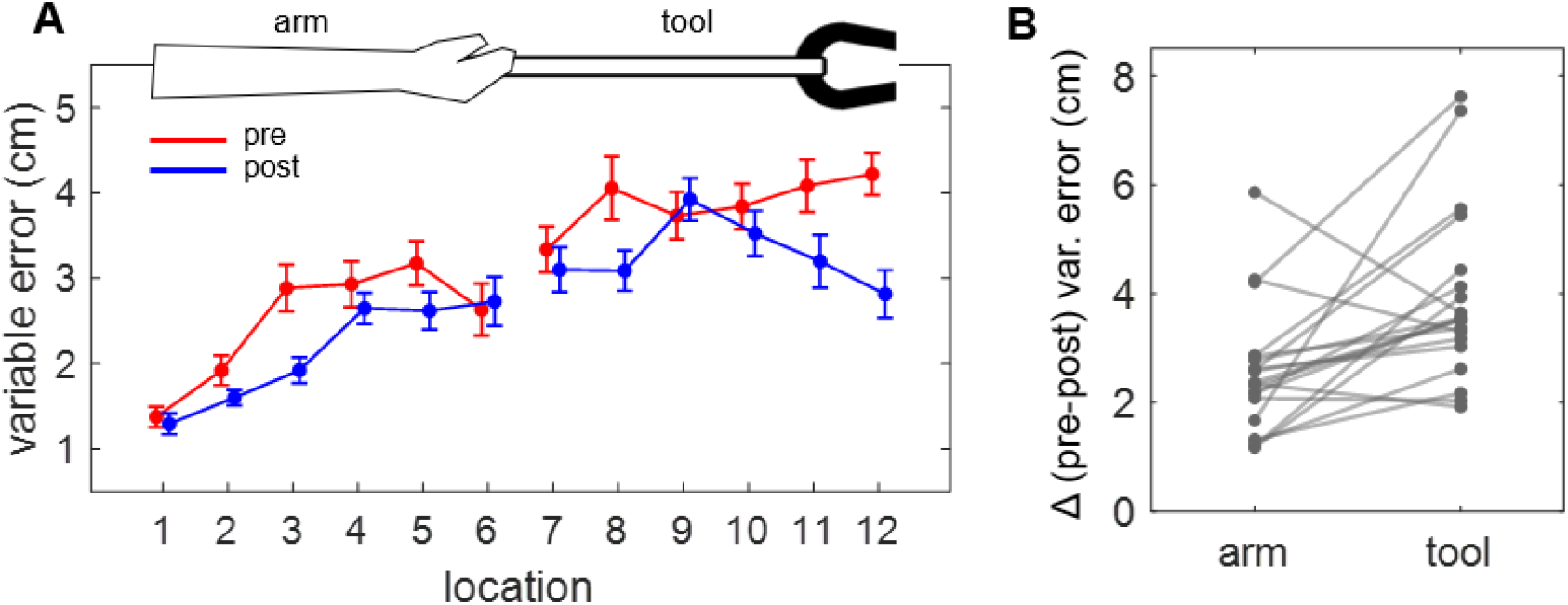
Variable errors in arm and tool mapping. **A**. The mean variable errors are plotted for each of the twelve target positions, for the pre tool use (red) and post tool use (blue). Error bars represent standard error of the mean. **B**. The six-dimensional Euclidean distances (Δ) computed on the variable errors between pre and post tool use are plotted for the arm and tool for each participant.

Pre-post changes in variable errors between arm and tool are not directly comparable with an ANOVA, since locations on the forearm and tool did not spatially correspond (i.e., arm location 1 was in a different spatial position compared to tool location 7). Therefore, to directly compare the magnitude of the tool-driven changes in variable error between arm and tool, we used Wilcoxon-signed rank test on the six-dimensional pre-post Euclidean distances. The test showed significant difference with z = -3.44, p = .003, effect size r = .770 (**Figure 3B**), indicating that variable errors for the tool decreased more from before to after tool use, compared to the arm.

## Discussion

Previous research has gathered abundant evidence on how healthy individuals are able to map the position of their body parts in space (e.g., Galigani et al., 2020; Longo & Haggard, 2010; Stone et al., 2021; Peviani et al., 2024); However, few studies have investigated if and how fine-grained spatial maps are created for tools that work as extensions of our limbs (Debats, Van De Langenberg, et al., 2010; Solomon & Burton, 1989; Burton & Turvey, 1990; Miller et al., 2018). In the present study, we aimed to investigate whether just holding a tool is sufficient for the sensorimotor system to readily map it in space, and whether such spatial map is comparable to that of the arm holding it. We used a spatial mapping task to measure the accuracy and precision of spatial estimates pertaining to several locations on the arm and tool. We then explored possible changes in map accuracy and precision after a period of tool use. We found that the spatial maps for the arm and hand-held tool were equally as accurate, with estimation errors that were comparable in direction and magnitude before and after tool use. While map accuracy did not change following tool use, we did observe a significant change in their spatial precision.

Whereas previous research has focused on the ability to perceive rod length via haptic feedback, our findings demonstrate the ability to perceive intermediate positions at fine-grained level. We know from previous research that participants are able to map tactile stimuli delivered on a hand-held rod (Miller et al., 2018; Miller et al., 2023), which provides indirect evidence that the sensorimotor system can infer a hand-held tool’s spatial features. Our work provides direct evidence that the sensorimotor system can build a spatial map of a tool, just by holding it. The somatic feedback gathered through mechanoreceptors of the hand and forearm while holding the tool (e.g., muscle torque, skin indentation) thus appears to be sufficient to build up a representation of the tool length that is comparable to that of the arm.

We did not find evidence of changes in the accuracy of the spatial maps following tool use. At first glance, this appears to be in contrast with previous research that observed changes in body representations following comparable tool use tasks (Canzoneri et al., 2013; Cardinali et al., 2009; Galigani et al., 2020; Miller et al., 2014). However, there are several differences between our experiment and those previous, which may contribute to our null result. A major difference between the present and previous research is that our paradigm required localizing several spatial locations rather than just the arm midpoint (i.e., arm bisection, Bruno et al., 2019; Romano et al., 2019; Sposito et al., 2012) or joints (Canzoneri et al., 2013; Cardinali et al., 2009; Galigani et al., 2020). Another difference from previous research is that participants, both before and after tool use, were required to perform spatial estimates of the arm *while* holding the tool. Under such conditions, the increased perceived arm length previously reported may not manifest since the tool was still physically present and working as a functional extension of the limb.

While the accuracy of spatial maps was unaltered following tool use, we found that the spatial resolution (i.e., precision) of the estimates improved for both the arm and the tool, with greater increase in precision for the tool compared to the arm. Importantly, increased precision was not spread uniformly across all tested locations but instead varied across them. This demonstrates that increased precision does not reflect a general effect of task repetition, rather a focal effect related to our experimental manipulation. Our plots (**Figure 3A**) and post-hoc tests show that the resolution of the spatial maps increased for central locations on the arm and for the distal boundary of the tool (i.e., tool tip). The close link between localization precision and somatosensory spatial computations (e.g., trilateration) (Miller et al., 2022; Fabio et al., 2024), suggests that tool use modified the computations that build the spatial maps of the arm and tool. We speculate that increased precision at the tool tip may reflect a more prominent role for the tip when computing location within the arm+tool system, following the extensive sensorimotor feedback gathered during the interaction with the tool. Future research involving computational modeling is needed to explore this possibility.

In all, our findings show that healthy individuals could map their arm and a hand-held tool in space with a comparable degree of accuracy, suggesting that the sensorimotor system can efficiently build up a spatial map of a tool based on available somatic feedback, such as skin stretch and torque. We also found that the spatial maps of the arm and tool did not change in terms of accuracy after tool use; rather, they changed in terms of precision, with a fine-tuning of the representation at key spatial landmarks, such as the tool tip. Our data thus clearly indicate that the sensorimotor system builds up a fine-grained spatial map of a tool as it does for the arm.

## Acknowledgments and fundings

V.C.P is supported by the Radboud Excellence Initiative grant. L.E.M is supported by an ERC 101076991 SOMATOGPS grant.

We extend our gratitude to the Brain Body Technology lab members for the scientific discussions, and the Technical Support Group of DCC for their assistance.

## Author contributions

V.C.P., L.E.M., G.R., and M.B. conceptualized the project. L.E.M., L.N.P, and V.C.P. designed the experiment. L.N.P collected the data. V.C.P. performed the formal analysis and visualized the data. V.C.P. and L.E.M. wrote the original draft of the manuscript. G.R. and M.B. provided feedback on the manuscript. L.E.M. supervised the project.

## Declaration of interests

The authors declare no competing interests.

## Notes

### Competing Interest Statement

The authors have declared no competing interest.

